# Model-based analysis of positive selection significantly expands the list of cancer driver genes, including RNA methyltransferases

**DOI:** 10.1101/366823

**Authors:** Siming Zhao, Jun Liu, Pranav Nanga, Yuwen Liu, A. Ercument Cicek, Nicholas Knoblauch, Chuan He, Matthew Stephens, Xin He

## Abstract

Identifying driver genes is a central problem in cancer biology, and many methods have been developed to identify driver genes from somatic mutation data. However, existing methods either lack explicit statistical models, or rely on very simple models that do not capture complex features in somatic mutations of driver genes. Here, we present driverMAPS (Model-based Analysis of Positive Selection), a more comprehensive model-based approach to driver gene identification. This new method explicitly models, at the single-base level, the effects of positive selection in cancer driver genes as well as highly heterogeneous background mutational process. Its selection model captures elevated mutation rates in functionally important sites using multiple external annotations, as well as spatial clustering of mutations. Its background mutation model accounts for both known covariates and unexplained local variation. Simulations under realistic evolutionary models demonstrate that driverMAPS greatly improves the power of driver gene detection over state-of-the-art approaches. Applying driverMAPS to TCGA data across 20 tumor types identified 159 new potential driver genes. Cross-referencing this list with data from external sources strongly supports these findings. The novel genes include the mRNA methytransferases METTL3-METTL14, and we experimentally validated METTL3 as a potential tumor suppressor gene in bladder cancer. Our results thus provide strong support to the emerging hypothesis that mRNA modification is an important biological process underlying tumorigenesis.

## Introduction

Cancer is caused by somatic mutations that confer a selective advantage to cells. Analyses of somatic mutation data from tumors can therefore help identify cancer-related (“driver”) genes, and this is a major motivation for recent large-scale cancer cohort sequencing projects^1^. Indeed, such analyses have already identified hundreds of driver genes across many cancer types^1,2^. Nonetheless, many important driver genes likely remain undiscovered^3^, especially in cancers with low sample sizes. Here we develop and apply new, more powerful, statistical methods to address this problem.

The basic idea underlying somatic mutation analyses is that genes exhibiting a high rate of somatic mutations are potential driver genes. However, mutation and repair processes are often significantly perturbed in cancer, so somatic mutations may also occur at a high rate in non-driver genes. Furthermore, somatic mutation rates vary substantially across genomic regions and across tumors. The challenge is to accurately distinguish driver genes against this complex background. Several main ideas have been developed to address this challenge. One idea is to carefully model the background somatic mutation process, by leveraging features that correlate with somatic mutation rate, such as replication timing^4^. Another idea is to utilize distinctive features of somatic mutations in driver genes: notably, mutations in driver genes tend to be more deleterious (“function bias”), and sometimes show a distinctive spatial pattern, tending to cluster together (e.g. in substrate binding sites)^5^. Methods that leverage one or more of these ideas include MuSiC^6^, MADGiC^7^, the Oncodrive suite^8–10^ and TUSON^11^.

Despite this progress, most existing methods do not explicitly model the process that generates the observed somatic mutations, namely, the interactions of mutational process and natural selection^12^. Indeed, tumorigenesis is well recognized as an evolutionary process^13,14^, and explicit modeling of mutation and selection is likely to be highly beneficial for analyzing somatic mutations in cancer^12, 15–17^. Many methods described above construct a null model for non-driver genes which lacks selection, and derive test statistics to reject this null model, without modeling of the alternative. Even recent, evolutionarily motivated models^16,17^ capture only the most basic impact of selection: differences in observed rates of nonsynonymous vs. synonymous mutations. Our approach, driverMAPS, is based on a much richer statistical model, which captures selection at the basepair level, and allows the strength of selection to depend on measures of functional importance such as conservation scores, SiFT^18^ and PolyPhen^19^. In addition, we use a Hidden Markov Model to capture potential spatial clustering of somatic mutations into “hotspots”. Our approach also introduces other innovative features: a detailed model of the background mutation processes, which accounts for known genomic features and variation across genes not captured by these features; and the use of a Bayesian hierarchical model to combine information across cancer types and hence improve parameter estimates.

Both simulations and application on TCGA data demonstrate the power of our approach. The explicit statistical models of driver and non-driver genes allow us to perform realistic simulations to assess methods, which was largely impossible in the past. We found that not all existing methods properly control the False Discovery Rate (FDR) for driver gene discovery, and among those with reasonable FDR control, driverMAPS has significantly higher power than existing ones. We applied driverMAPS to TCGA exome sequencing data from 20 cancer types. The results suggest that driverMAPS is better able to detect previously known driver genes than existing methods, without excessive false positives. In addition, driverMAPS identified 159 new potential driver genes not identified by other methods. Both literature survey and extensive computational validation suggest that many of these genes are likely to be true driver genes. The novel potential driver genes included both METTL3 and METTL14, which together form a key enzyme for RNA methylation. We experimentally validated the functional relevance of somatic mutations in METTL3, providing further support for both the effectiveness of our method, and for the potential importance of RNA methylation in cancer. We believe that our methods and results will facilitate the future discovery and validation of many more driver genes from cancer sequencing data.

## Results

### driverMAPS: a probabilistic model of somatic mutation selection patterns

Our approach is outlined in Figure 1. In brief, we model aggregated exonic somatic mutation counts from many tumor samples (e.g. as obtained from a normal-tumor paired sequencing cohort). Let Y_g_ denote the mutation count data in gene g. We develop models for Y_g_ under three different hypotheses: that the gene is a “non-driver gene” (H_0_), an “oncogene” (H_OG_) or a “tumor suppressor gene” (H_TSG_). Each model has two parts, a background mutation model (BMM), which models the background mutation process, and a selection mutation model (SMM), which models how selection acts on functional mutations. The rate of observed mutation at a position is the product of the background mutation rate (from BMM) and a coefficient reflecting the effect of position-specific selection (from SMM). We note that the coefficient can be related to the selection coefficient of the mutation and effective population size under a simplified population genetic model^12^. If the coefficient is greater than 1, it indicates positive selection and if it is less than 1, negative selection. The BMM parameters are shared by all three hypotheses, reflecting the assumption that background mutation processes are the same for cancer driver and non-driver genes. In contrast the SMM parameters are hypothesis-specific, to capture the different selection pressures in oncogenes vs tumor suppressor genes vs non-driver genes. We fit the hypothesis-specific parameters using training sets of known oncogenes^1^ (H_OG_), known TSGs^1^ (H_TSG_), and all other genes (H_0_). (This last set will contain some -- as yet unidentified -- driver genes, which will tend to make our methods conservative in terms of identifying new driver genes.) To combine information across tumor types we first estimate parameters separately in each tumor type, and then stabilize these estimates using Empirical Bayes shrinkage^20^.

**Figure 1.**
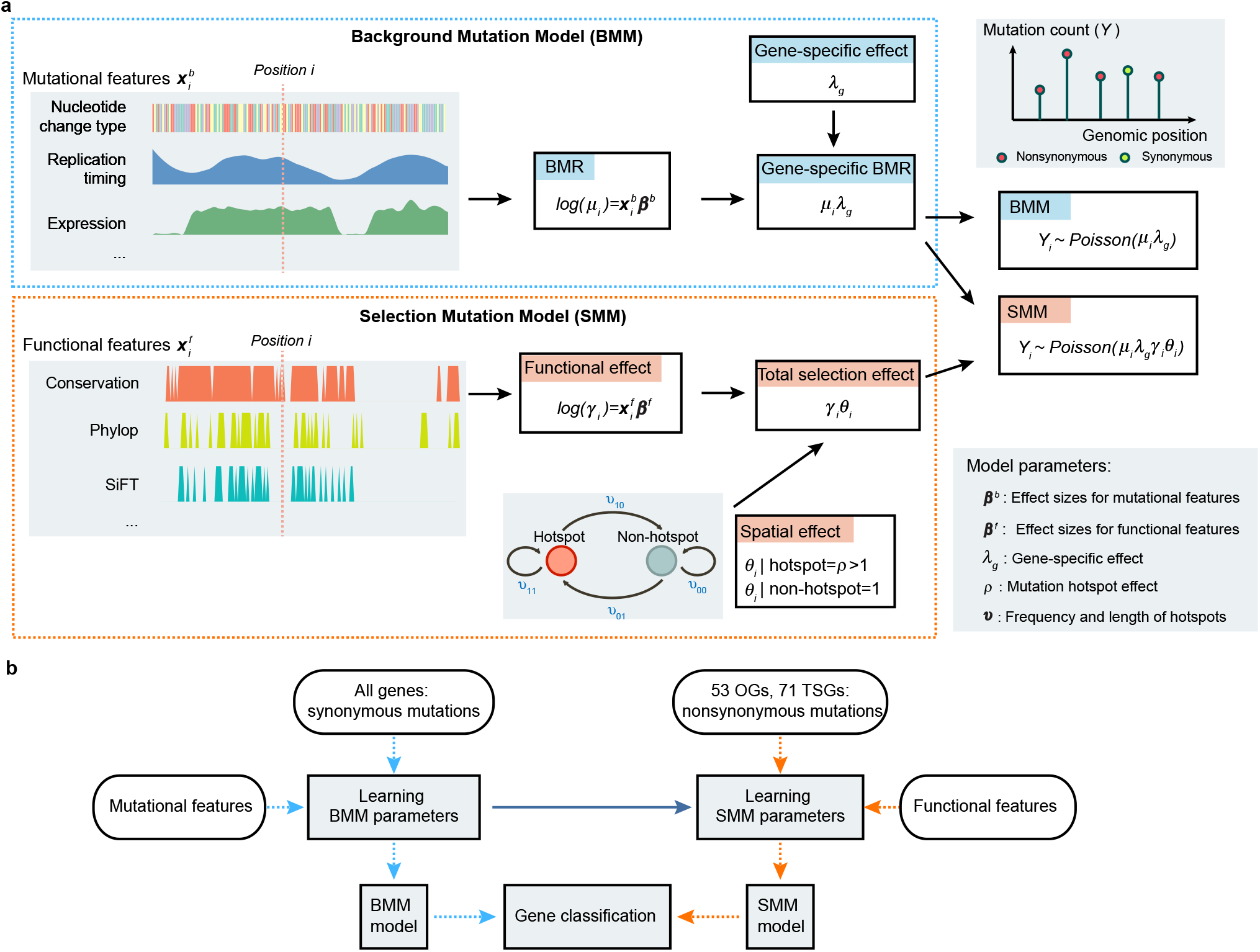
Overview of the model-based framework driverMAPS for cancer driver gene discovery. **(a)** Base-level Bayesian statistical modeling of mutation count data in driverMAPS. For positions without selection, the observed mutation rate is modeled by Background Mutation Model (BMM). Under BMM, the Background Mutation Rate (BMR)(*μ_i_*) is determined by the log-linear model that takes into account known mutational features and further adjusted by gene-specific effect (*λ_g_*) to get gene-specific BMR (*μ_i_ λ_g_*). For positions under selection, the observed mutation rate is modeled as gene-specific BMR adjusted by selection effect (Selection Mutation Model, SMM). The selection effect has two components: functional effect (*γ_i_*) takes into account functional features of the position by the log-linear model and spatial effect (*θ_i_*) takes into account the spatial pattern of mutations by Hidden Markov Model. For both BMM and SMM, given the mutation rate, the observed mutation count data is modeled by Poisson distribution. Note: we simplify the model to only show mutation rate at position *i*, ignoring allele specific effect for illustration purposes. See Methods for full parameterization. (**b)** Gene classification workflow. Parameters in BMM are estimated using synonymous mutations from all genes. This set of parameters is fixed when inferring parameters in SMM. To infer parameters in SMM, we use nonsynonymous mutations from known OGs or TSGs. driverMAPS then performs model selection by computing gene-level Bayes Factors to prioritize cancer genes.

Having fit these models, we use them to identify genes whose mutation data are most consistent with the driver genes models (H_OG_ and H_TSG_). Specifically, for each gene g, we measure the overall evidence for g to be a driver gene by the Bayes Factor (likelihood ratio), BF_g_, defined as:

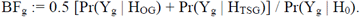

Large values of BF_g_ indicate strong evidence for g being a driver gene, and at any given threshold we can estimate the Bayesian FDR. For results reported here we chose the threshold by requiring FDR<0.1.

### driverMAPS effectively captures factors influencing somatic mutations

We used a total of 734,754 somatic mutations from 20 tumor types in the TCGA project as our input data^21^. We focused on single nucleotide somatic variations and extensively filtered input mutation lists to ensure data quality (see Methods). Figure S1 summarizes mutation counts and cohort sizes.

The first step of our method is to estimate parameters of the Background Mutation Model (BMM) using data on synonymous mutations. These parameters capture how mutation rates depend on various “background features” (Table S1), which include mutation type (C>T, A>G, *etc*), CpG dinucleotide context, expression level, replication timing and chromatin conformation (HiC sequencing)^4^. The signs and values of estimated parameters were generally similar across tumor types, and consistent with previous evidence for each feature’s effect on somatic mutation rate. For example, the estimated effect of the feature “expression level” was negative for almost all tumors, consistent with transcriptional coupled repair mechanisms effectively reducing mutation rate (Figure S2).

Our BMM also estimates gene-specific effects, using synonymous mutations of a gene, to allow for local variation in somatic mutation rate not captured by measured features. Intuitively, the gene-specific effect adjusts a gene’s estimated mutation rate downward if the gene has fewer synonymous mutations than expected based on its known features, and upwards if it has more synonymous mutations than expected. A challenge here is that the small number of mutations per gene (particularly in small genes) could make these estimates inaccurate. Here we address this using Empirical Bayes methods to improve accuracy, and avoid outlying estimates at short genes that have few potential synonymous mutations (Figure 2a). Effectively, this adjusts a gene’s rate only when the gene provides sufficient information to do so reliably (sufficiently many potential synonymous mutations). To demonstrate the reliability of the resulting estimates we use a procedure similar to cross-validation: we estimated each gene’s gene-specific effect using its synonymous mutations, and then test the accuracy of the estimate (compared to no gene-specific effect) in predicting the number of nonsynonymous mutations. We assume that for the vast majority of genes, their mutational counts are dominated by background mutation processes, rather than selection. Figure 2b shows results for SKCM tumors: without gene-specific effect the correlation of observed and expected number of nonsynonymous mutations across genes was 0.56; with gene-specific adjustment the correlation increased to 0.88. Similar improvements were seen for other tumors (Figure S3).

**Figure 2.**
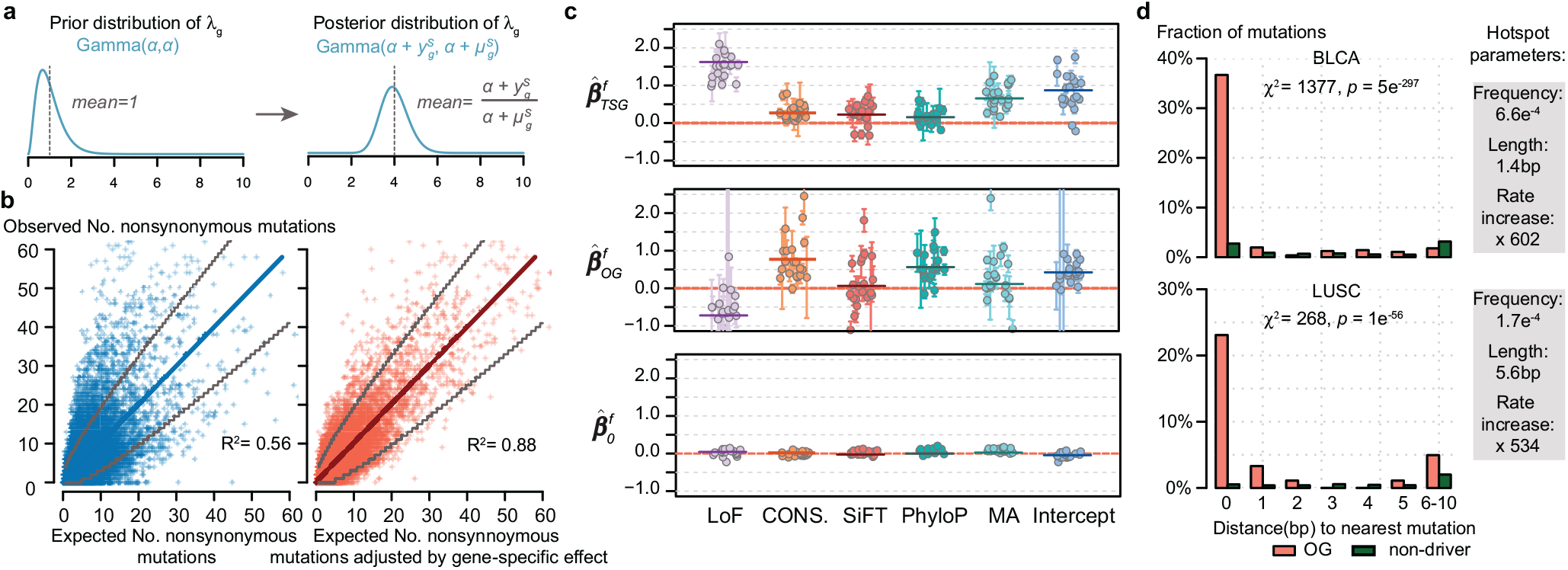
Parameter estimation results for gene-specific, functional and spatial effects. **(a)** Schematic representation of how fitting synonymous mutation data affects estimation of gene-specific effect (*λ_g_*). Note the difference between the prior and posterior distributions of *λ_g_*. α is a hyperparameter, 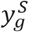 and 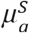 are the observed and expected number of synonymous mutations in gene *g*, respectively. **(b)** Improved fitting of observed number of nonsynonymous mutations in genes with gene-specific effect adjustment. Data from tumor type SKCM was used. The adjustment here is the posterior mean of λ_g_ fitting synonymous mutation data 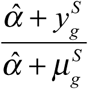. Each dot represents one gene. Grey lines indicate upper and lower bounds of 99% confidence interval from Poisson test. The diagonal line has slope =1 and R^2^ was calculated using this as the regression line. **(c)** Effect sizes for five functional features and average increased mutation rate for TSGs (top), OGs (middle) and non-driver genes (bottom). Each dot represents an estimate from one tumor type. Horizontal bars represent mean values after shrinkage. All features are binarily coded. LoF, loss-of-function (nonsense or splice site) mutations or not. CONS., amino acid conservation; SiFT, PhyloP and MA, predictions from software SiFT^18^, PhyloP^43^ and MutationAssessor^44^, respectively; intercept, average increased mutation rate. **(d)** Fraction of mutations that has the nearest mutation 0,1,2,.. bp away, where 0bp means recurrent mutations. Data from tumor type BLCA and LUSC was used. The test statistics *χ*^2^ and *p* values were obtained in the spatial model selection procedure (see method, Table S6). Inferred parameters related to the spatial model are shown on the right.

The next step is to estimate parameters of the Selection Mutation Models (SMM), using data on non-synonymous mutations. These parameters capture how the rate of non-synonymous somatic mutations depend on various “functional features” (Table S2-S4), including loss-of-function (LoF) status, conservation scores, *etc*. Signs and values of estimated parameters were generally similar across tumor types, and consistent with their expected impact on gene function (Figure 2c). For example, the estimated effect of the “LoF” feature was positive for H_TSG_ and negative for H_OG_, indicating that loss-of-function mutations are enriched in TSGs and depleted in OGs, as expected from their respective roles in cancer. The intercept terms for both TSG and OG are positive, suggesting that somatic mutations are enriched in both types of cancer driver genes.

The final step is to estimate parameters of the spatial model (HMM, Figure 1), which are designed to capture how somatic mutations may cluster together in “hotspots” in driver genes. Preliminary investigations showed that spatial clustering is generally stronger in known OGs than in known TSGs, and so we fit the spatial model separately for OGs and TSGs in each tumor type (Table S5). Our model identified some tumor types (e.g. BLCA and LUSC, Figure 2d) with strong spatial clustering. In BLCA, the estimated hotspots are very short (mean 1.4bp) and are primarily capturing an excess in recurrent mutations (independent mutations at the same base) compared with expectations (Figure 2d). In LUSC, the clustering extends over slightly longer regions (mean 5.6bp), but still the primary signal is an excess of recurrent mutations (Figure 2d).

### Simulations demonstrate that driverMAPS improves detection of driver genes

While many methods have been developed for driver gene identification, it is difficult to compare them on real data where the true status of each gene is often unknown. Simulations are extremely valuable in such situations, and have been used in many fields, including population genetics^22^, statistical genetics^23^ and single-cell transcriptomics^24^. Here we exploit our explicit statistical model to perform realistic simulations based on parameters inferred from real data (here, the TCGA UCS cohort).

We first assess a common strategy used in the field: Fisher’s method to combine p-values of a gene, each capturing a single feature of positive selection. We simulated somatic mutations in a positively selected gene with both increased nonsynonymous mutation rates and mutational hotspots. We ran two simple tests -- a dN/dS test to detect enrichment of functional mutations and another to detect spatial clustering (see Methods) -- and then combined *p* values using Fisher’s method. Perhaps unexpectedly, the combined test has lower power than the dN/dS test alone (Figure 3a). We believe that this is because spatial clustering is a relatively weak feature in our simulations (as in real data) and so the spatial test has much less power than the dN/dS test. Consequently the spatial test adds more noise than signal, decreasing power. This result highlights a weakness of methods based on combining *p* values; model-based approaches, such as ours, avoid this problem by automatically weighting different features of the data based on their informativeness.

**Figure 3.**
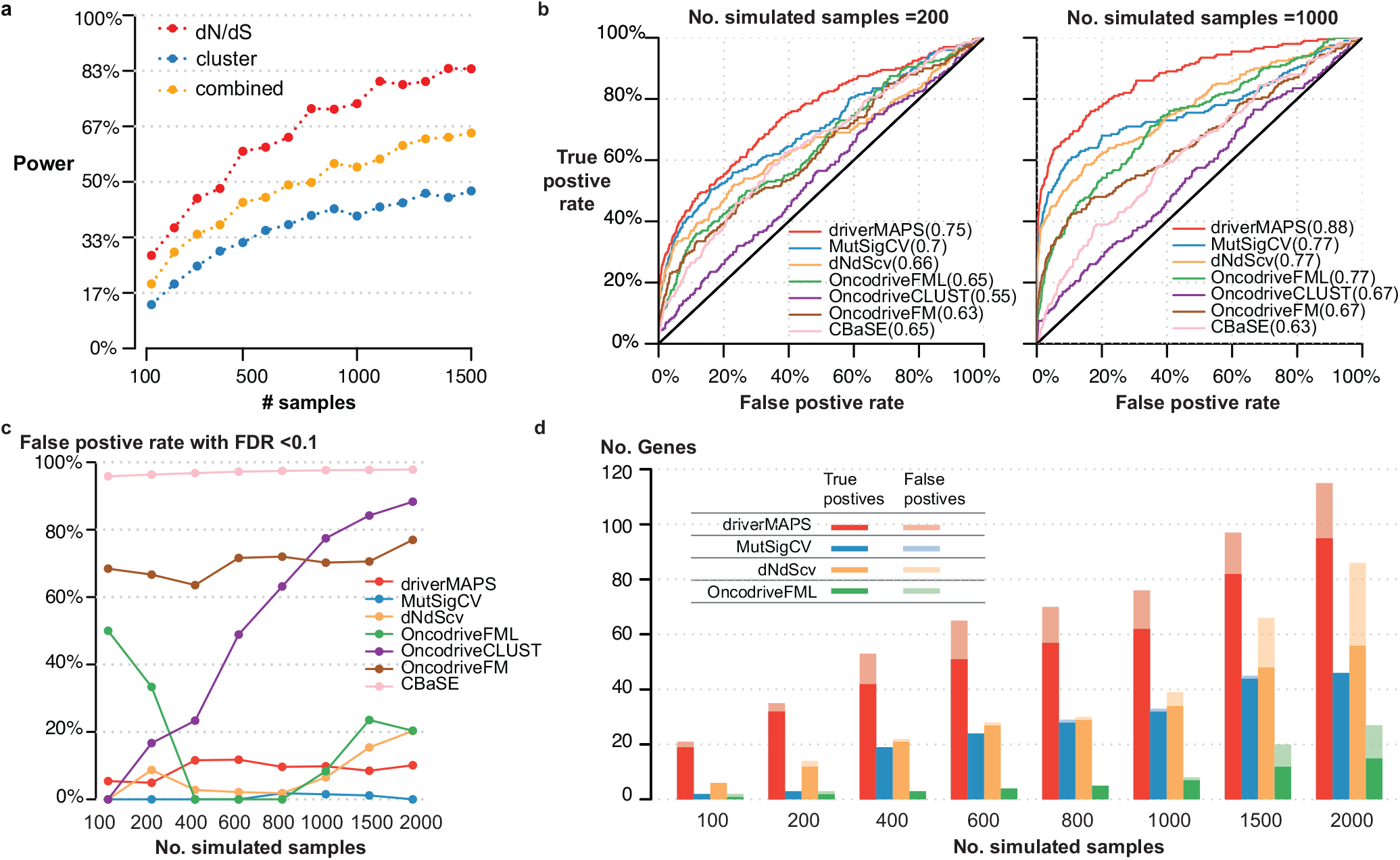
driverMAPS predicts driver genes with high accuracy and increased power in simulations. **(a)** Combining p values from methods that use only one feature of positive selection a time will lose power. We simulated mutations of a gene under positive selection under various sample sizes, then assessed the power of detecting this gene as positively selected. “dN/dS” only captures the excess of nonsynonymous mutations, “cluster” only captures spatial clustering pattern of mutation, “combined” combines p values from “dN/dS” and “cluster” using Fisher’s method. **(b)** Receiver Operating Characteristic (ROC) curves of several methods applied to genome-wide simulation data. 324 genes are chosen to be positively selected (191 TSGs and 133 OGs) and the rest of genes are neutral. We used 124 out of the 324 genes as training set for driverMAPS and used the rest 200 genes as the test set to generate ROC curves. Area Under an ROC Curve (AUROC) values are shown in parentheses. **(c)** False positive rate at FDR cutoff 0.1 on the simulated data. **(d)** Number of true positive and false positive genes at FDR<0.1. We did not count the 124 training genes (for driverMAPS) to ensure a fair comparison among methods.

We next used simulations to compare driverMAPS with six existing algorithms: MutSigCV, OncodriveFML^9^, OncodriveFM^10^, OncodriveCLUST^8^, dNdScv^16^ and CBaSE^17^. We performed simulations of all genes in the genome where 324 genes are randomly chosen as oncogenes or tumor suppressor genes. We found that, for distinguishing driver vs non-driver genes, driverMAPS outperformed all other methods (Figure 3b). Furthermore, only driverMAPS and MutSigCV consistently control FDR across all sample sizes (Figure 3c). Excluding three methods with obvious problems of FDR control (OncodriveFM, OncodriveCLUST, CBaSE), driverMAPS identifies the most driver genes at FDR < 0.1 (Figure 3d). Overall we found the power of driverMAPS to discover novel driver genes can be double that of other leading methods (and even more in smaller samples).

### Application of driverMAPS on TCGA data

We next compared results from driverMAPS and other algorithms for predicting driver gene using TCGA data (see Methods). Besides the full implementation of driverMAPS, we also tried a “basic” version that looks only for an excess of nonsynonymous somatic mutations (without any functional features or spatial model), and a “+feature” version with functional features but not the spatial model. We applied all methods to the same somatic mutation data and compared the genes they identified with a list of “known driver genes” (713 genes) compiled as the union of COSMIC CGC list (version 76)^25^, Pan-Cancer project driver gene list^2^ and list from Vogelstein B (2013)^1^ (see Supplementary Note). To avoid overfitting of driverMAPS to the training data, we trained driverMAPS with a leave-one-out strategy in these assessments.

For each method we computed both the total number of genes detected (at FDR=0.1) (Figure 4a) and the “precision” -- the fraction that are on the list of known driver genes (Figure 4b). All versions of driverMAPS identified more driver genes than either MutSigCV, dNdScv or OncodriveFML, while maintaining a similarly high precision. The full version of driverMAPS (with the spatial and functional features) identified nearly twice more genes. Furthermore, this higher detection rate of driverMAPS was consistent across tumor types (Figure 4c). The other methods, OncodriveFM, OncodriveCLUST and CBaSE, behaved quite differently, identifying thousands of driver genes but with much lower precision, consistent with poor FDR control in simulations (Figure 3c). For OncodriveFM and OncodriveCLUST, the lowest precision was in the tumor types with the highest mutation rates (e.g. BLCA, LUSC, LUAD), suggesting the accuracy of these methods may be affected by mutation rates (Figure S4). While precision of OncodriveFM and OncodriveCLUST showed a negative correlation with mutation rate (Pearson r = −0.44 and −0.56), the precision of driverMAPS showed negligible correlation (Pearson r = 0.05).

**Figure 4.**
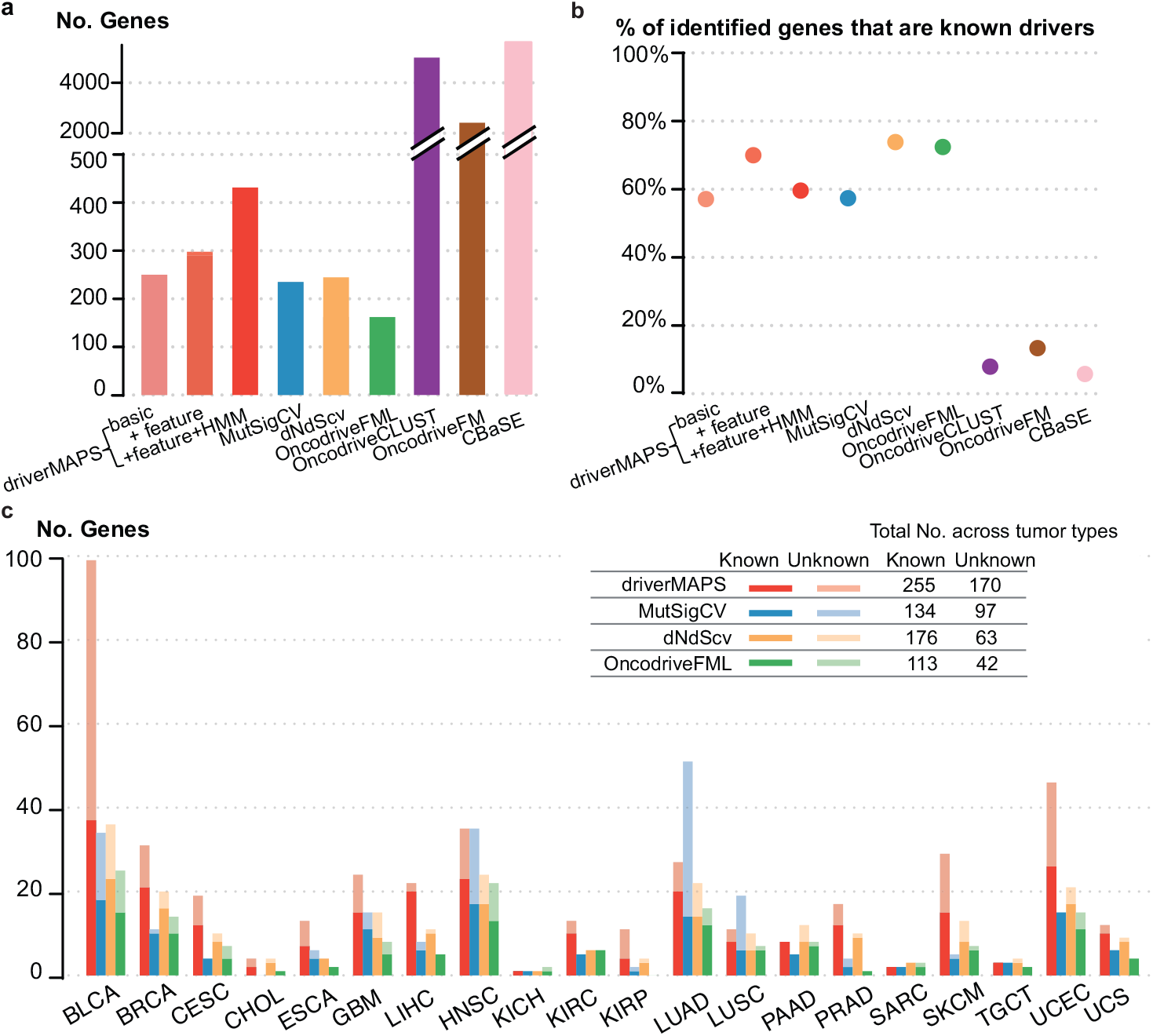
Gene prediction using TCGA somatic mutation data. **(a)** Total number of predicted driver genes aggregating across all cancer types. driverMAPS (Basic), driverMAPS with no functional features information and no modeling of spatial pattern; driverMAPS (+ feature), driverMAPS with all five functional features in Figure 2, no modeling of spatial pattern; driverMAPS (+feature + HMM), complete version of driverMAPS with all five functional features and spatial pattern. **(b)** Percentage of known cancer genes among predicted driver genes aggregating across all cancer types. **(c)** Number of significant genes at FDR<0.1 stratified by tumor type. For all “Unknown” genes included here, we verified mutations by visual inspection of aligned reads using files from Genomic Data Commons (see Supplementary notes). Total numbers of known and unknown significant genes aggregating across all cancer types are summarized topright.

### Evaluation of potential novel drivers identified by driverMAPS

Summing across all 20 tumor types, at FDR 0.1, driverMAPS identified 255 known driver genes and 170 putatively novel driver genes (159 unique genes across the 20 tumor types; 70 classified as TSGs and 100 as OGs; Figure 5a, Table S7). Almost half of these putative novel genes were not called by MutSigCV, OncodriveFML or dNdScv. Ten novel genes were found independently in at least two tumor types (Table 1). This is unlikely to happen by chance (permutation test, *p* < 1e^-4^), so these genes seem especially good candidates for being genuine driver genes.

**Figure 5.**
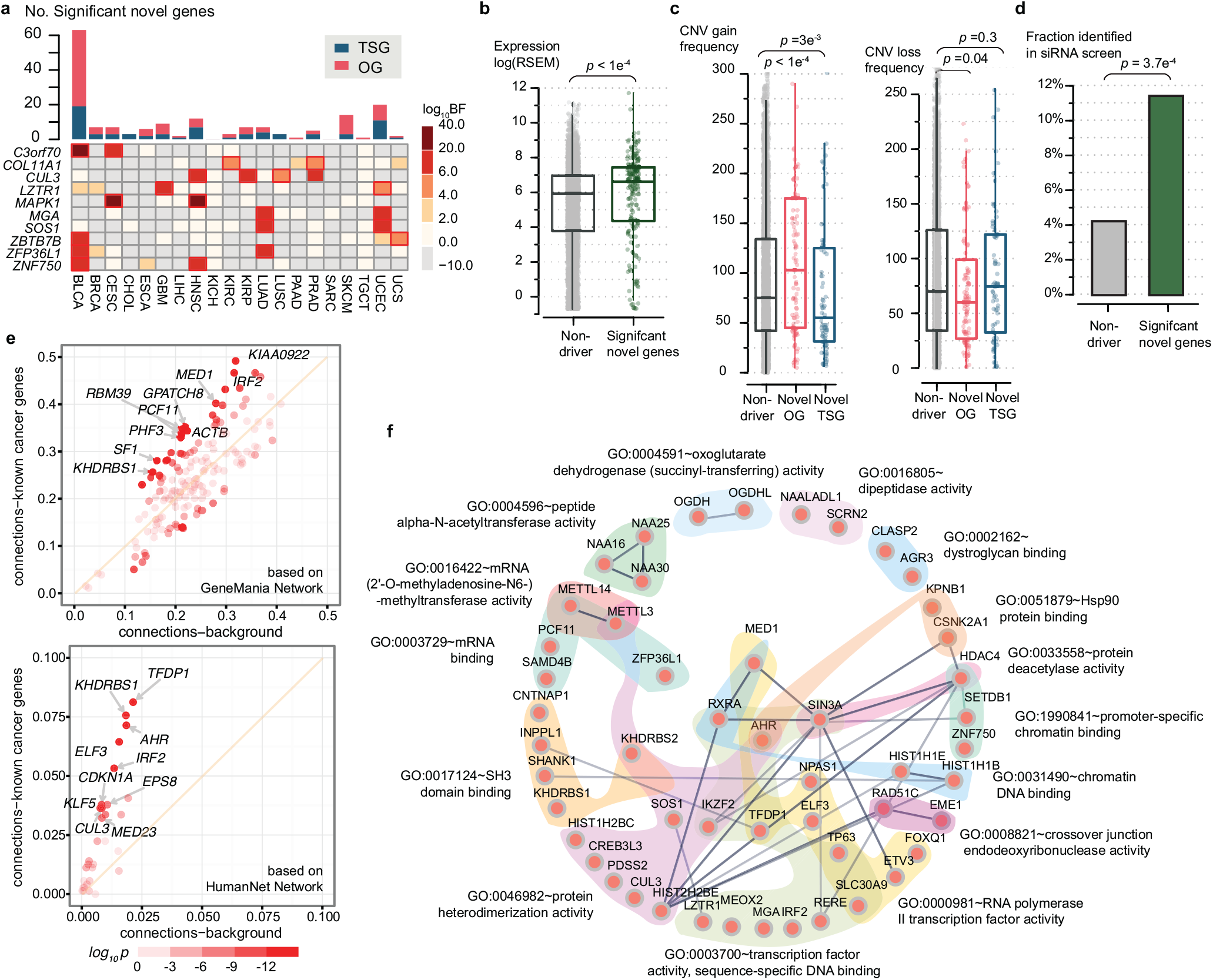
Evaluation of novel cancer genes predicted by driverMAPS. **(a)** Overview of predicted novel cancer genes. Top, number of novel genes in each cancer type. Bottom heatmap of Bayes factors (BF) for recurrent novel genes across tumor types. Significant Bayes factors are highlighted by red boxes. **(b-d)** Predicted novel cancer genes show known cancer gene features. For each feature, quantification of the feature level in the novel cancer gene set was compared to the non-drive (neither known or predicted) gene set. The features are gene expression levels^21^ stratified by tumor type the novel genes were identified from (b), similarly stratified copy number gain/loss frequencies^21^ (c) an fraction of genes identified in a siRNA screen study^26^ (d). In (b) and (c), the center line, median; box limits, upper and lower quartiles. **(e)** Enriched connectivity of a predicted gene with 713 known cancer genes (Y-axis) compared to with all genes (n=19,512, X-axis). Connectivity of a selected gene with a gen set is defined as the number of connections between the gene and gene set found in a network databas divided by the size of the gene set. Each dot represents one of the 159 novel genes with 10 most enriched ones labeled. Color of dots indicates two-sided Fisher exact *p* value for enrichment. **(f)** Significantly enriched GO-term gene sets (FDR < 0.1, “molecular function” domain) in predicted novel cancer gene GO-term^29,30^ gene sets are indicated by distinct background colors. Links among genes represent interaction based on STRING network database^45^ with darker color indicating stronger evidence.

**Table 1.** Novel significant drivers found in at least two tissue types

| Gene | #Missense | #LoF | #Silent | log<sub>10</sub>BF | Tumor | Function |
| --- | --- | --- | --- | --- | --- | --- |
| **C3orf70** | 14/3 | 1/1 | 0/0 | 9.3/3.8 | BLCA/CESC | Unknown |
| **COL11A1** | 7/13 | 4/2 | 0/0 | 2.2/2.2 | KIRC/PRAD | Collagen formation, expression associated with colorectal, ovarian cancers, etc (23934190, 11375892) |
| **CUL3** | 15/8/4 | 5/4/0 | 1/0/0 | 3.5/3.8/2.6 | HNSC/KIRP/PRAD | Core component of E3 ubiquitin ligase complex, with many downstream targets affecting carcinogenesis, like NRF2 (24142871) |
| **LZTR1** | 9/10 | 0/1 | 0/2 | 2.9/2.1 | GBM/UCEC | Adaptor of CUL3-containing E3 ligase complexes, inactivation drives glioma self renewal and growth (23917401) |
| **MAPK1** | 9/7 | 0/1 | 0/0 | 15.1/12.8 | CESC/HNSC | MAP kinase. The MAPK/ERK cascade has important well characterized and important roles in cancer (17496922) |
| **MGA** | 35/11 | 16/5 | 5/3 | 3.8/2.7 | LUAD/UCEC | Dual-specificity transcription factor, can inhibit MYC-dependent cell transformation (10601024) |
| **SOS1** | 12/7 | 1/0 | 3/0 | 3.5/7.0 | LUAD/UCEC | Guanine nucleotide exchange factor for RAS proteins, which are well-known for roles in cell proliferation (17486115) |
| **ZBTB7B** | 11/5 | 1/1 | 0/0 | 6.2/2.3 | BLCA/UCS | Transcriptional regulator of lineage commitment of immature T-cell precursors (17878336) |
| **ZFP36L1** | 12/11 | 4/3 | 1/0 | 3.4/5.2 | BLCA/LUAD | Involved in mRNA degradation. Deletion leads to T lymphoblastic leukemia (20622884) |
| **ZNF750** | 17/13 | 3/7 | 2/1 | 3.4/5.1 | BLCA/HNSC | An essential regulator of epidermal differentiation. Depletion promotes cell proliferation in ESCA (24686850) |

We use “/” to separate data obtained from different tumor types as indicated in the “Tumor” column. A brief description of the gene’s function and its known role in cancer is provided in the “Function” column. Reference PMIDs are given in parentheses.

Since it is impractical to functionally validate all 170 putative novel genes, we sought other data to support these genes likely being involved in cancer. We first selected three common cancers -- breast, lung and prostate -- and conducted an extensive literature survey for each novel gene identified in these tumor types. Among a total of 22 novel genes, we found clear support in the literature for 20 being involved with cancer biology, either directly implicated as oncogenes or tumor suppressor genes (but not in the list of “known driver genes”) or linked to well-established cancer pathways (Table S8).

We next assessed whether the novel genes were enriched for features often associated with driver genes. Previous studies reported that driver genes tend to be highly expressed^4^ compared with other genes, and indeed we found that, collectively, the novel genes showed significantly higher expression than randomly sampled genes in the corresponding tissues^21^ (*p*<1e^-4^) (Figure 5b).

Previous studies have also reported that driver genes tend to show enrichment and depletion for different copy-number-variation (CNV) events, depending on their role in cancer. Specifically, OGs are enriched for CNV gains and depleted for CNV loss, whereas TSGs show enrichment for loss and depletion for gains. Consistent with this, we found novel genes identified as OGs are enriched for CNV gain events (*p*<1e^-4^) while novel TSGs are depleted (*p*=3e^-3^). CNV loss events for novel OGs are depleted compared to novel TSGs and to other genes (*p*= 0.04) (Figure 5c).

We also compared our novel genes with a “cancer dependency map” of 769 genes identified from a large-scale RNAi screening study across 501 human cancer cell lines^26^. These are genes whose knockdown affects cell growth differently across cancer cell lines, thus likely representing genes that are critical for tumorigenesis, but not universally essential genes. We found 16 novel driver genes overlapped with this gene list, a significant enrichment compared with random sampling (odds ratio 2.9, *p*=3.7e^-4^) (Figure 5d and Table S9).

To test whether our novel genes are functionally related to known cancer driver genes we examined the connectivity of these two sets of genes in the HumanNet^27^ gene network, which is built from multiple data sources including protein-protein interactions and gene co-expression. On average, each novel gene is connected to 3.8 known driver genes, significantly higher than expected by chance (*p* = 0.001). We obtained a similarly significant result using a different gene network, GeneMania^28^, which is constructed primarily from co-expression (*p* = 0.008) (Figure 5e).

Finally, we identified enriched functional categories in our novel genes using GO enrichment^29,30^ analysis (by geneSCF^31^). Significant GO terms (FDR < 0.1, Figure 5f) include many molecular processes directly implicated in cancer, such as transcription initiation and regulation. The significant terms also include several that have not been previously implicated in cancer. Genes NAA25, NAA16 and NAA30 (GO: 0004596) are peptide N-terminal amino acid acetyltransferases^32^. NATs are dysregulated in many types of cancer, and knockdown of the NatC complex (NAA12-NAA30) leads to p53-dependent apoptosis in colon and uterine cell lines^33^. OGDH and OGDHL (GO:0004591) have oxoglutarate dehydrogenase activities and part of the tricarboxylic acid (TCA) cycle^34^. METTL3 and METTL14 (GO: 0016422) form the heterodimer N6-methyltransferase complex, and are responsible for methylation of mRNA (m^6^A modification)^35^. This form of RNA modification may influence RNA stability, export and translation, and has been shown to be important for important biological processes such as stem cell differentiation. Our results suggest that this RNA methylation pathway may also play a key role in tumorigenesis, and so we examined the results for these genes in more detail.

### METTL3 is a potential TSG in bladder cancer

driverMAPS identified the genes METTL3 and METTL14 as driver genes in the cohorts BLCA (bladder cancer) and UCEC (uterine cancer) respectively. These two genes had relatively low mutation frequencies (4% and 2%) and were not detected by MutSigCV, dNdScv or OncodriveFML (those with reasonable FDR control). Inspecting the mutations in these two genes, we found many to be “functional” as predicted by annotations, and showed spatial clustering patterns in the MTase domain (Figure 6a). Furthermore METTL3 contained a single synonymous mutation, and METTL14 contained none, suggesting low baseline mutation rates at the two genes. While this manuscript was in preparation, METTL14 was independently identified as a novel TSG in endometrial cancer (Chuan He, to appear). We thus focused on METTL3 in bladder cancer.

**Figure 6.**
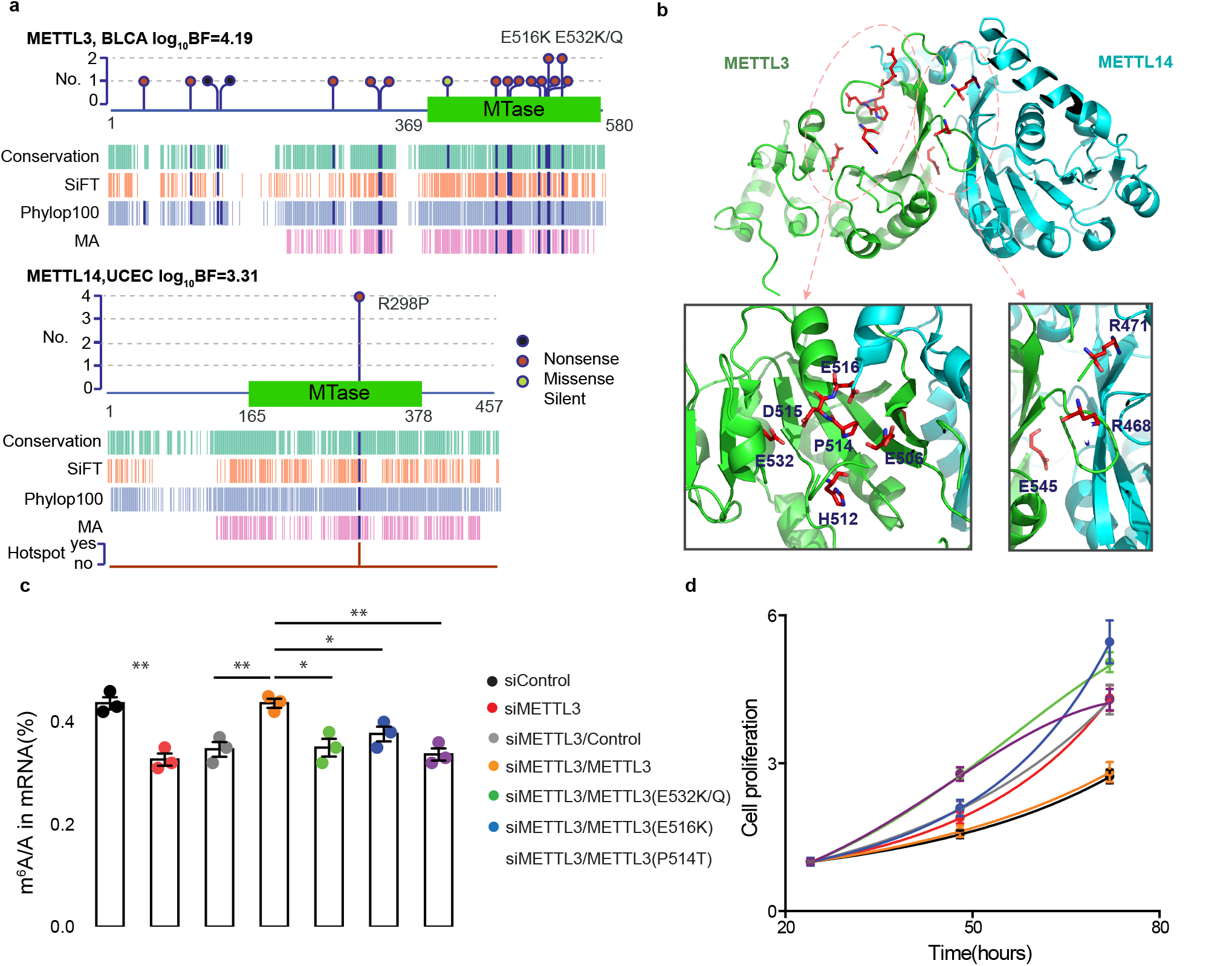
Functional validation of METTL3 as a TSG in bladder cancer. **(a)** Features of mutations in METTL3 and its heterodimerization partner METTL14. We show schematic representations of protein domain information and mark mutation positions by “lollipops”. Recurrent mutations are labeled above. Start and end of domain residues are labeled below. Dark blue bars in aligned annotation tracks indicate the mutation is predicted as “functional”. Track “Hotspot” is the indicator of whether the mutation is in hotspot or not in driverMAPS’s spatial effect model (See supplementary note). **(b)** Structural context of METTL3 mutations revealed two regional clusters. Top, structure of METTL3 (residues 369–570) and METTL14 (residues 117–402) complex (PDB ID: 5IL0) with mutated residues in stick presentation. Bottom, zoom-in views of the two regions with mutated residues labeled. **(c)** Impaired m^6^A RNA methyltransferase activity of mutant METTL3 in bladder cancer cell line “5637”. LC-MS/MS quantification of the m^6^A/A ratio in polyA-RNA in METTL3 or Control knockdown cells, rescued by overexpression of wildtype or mutant METTL3 is shown. **(d)** Mutant METTL3 decreased proliferation of “5637” cells. Proliferation of METTL3 or Control knockdown cells, rescued by overexpression of wildtype or mutant METTL3 in MTS assays is shown. Cell proliferation is calculated as the MTS signal at the tested time point normalized to the MTS signal ~ 24 hours after cell seeding. For all experiments in **(c-d)**, number of biological replicates is 3 and error bars indicate mean ± s.e.m. *, *p* < 0.05; **, *p* < 0.01 by two sided *t*-test. Legend is shared between (c) and (d).

To gain further insights into the potential impact of the somatic mutations in METTL3, we performed structural analysis. By mapping mutations in the MTase domain of METTL3 to its crystal structure^36^, we found them to be concentrated in two regions: one close to the binding site of S-Adenosyl methionine (AdoMet, donor of the methyl group) and the other in the putative RNA binding groove at the interface between METTL3 and METTL14 (Figure 6b). The region close to the AdoMet binding site contains seven mutations: E532K, E532Q, E516K, D515Y, P514T, H512Q and E506K. Position E532 has been reported to form direct water-mediated interactions with AdoMet^36^. The other mutations map to gate loop 2 (E506K and E516K map to the start and end; the other three mutations are inside the loop) which is known to undergo significant conformational change before and after AdoMet binding. Thus all these mutations are good candidates for affecting adenosine recognition. The second region, in the METTL3-METTL14 interface, contains mutations R471H, R468Q and E454K, and so these mutations are good candidates for disrupting METTL3-METTL14 interaction. In further support of this, the highly recurrent R298P mutation in METTL14 lies in the binding groove of the METTL14 gene.

We performed functional experiments to test whether mutations (n=7) in the first region affect METTL3 function. In an *in vitro* assay, most mutations reduced methyltransferase activity of METTL3 (Figure S5, see methods) and we chose four mutations (at three positions) for further cell line experiments. In two bladder cell lines (“5637” and “T24”), knock down of METTL3 by siRNA significantly reduced m6A methyltransferase activity (Figure 6c for “5637”, Figure S6a for “T24”). When we tried to rescue this phenotype by transfection of METTL3 mutants, all of the mutations, E532K/Q, E516K and P514T failed to restore methyltransferase activity to original levels (Figure 6c, Figure S6a), suggesting that they are loss-of-function mutations.

We next examined whether disruption of METTL3 is associated with tumor progression. Indeed, knockdown of METTL3 significantly increased cell proliferation. Wild type METTL3 successfully restored the cells to their normal growth rate but none of the mutants could (Figure 6d, Figure S6b).

These results show that somatic mutations in METTL3 may promote cancer cell growth by disrupting the RNA methylation process, and invite further characterization of the role of METTL3 and RNA methylation in tumorigenesis by in vivo experiments.

## Discussion

We have developed an integrated statistical model-based method, driverMAPS, to identify driver genes from patterns of somatic mutation. By applying this method to data from multiple tumor types from TCGA, we detected 159 novel potential driver genes. We experimentally validated the function of mutations in one gene, METTL3. The remaining genes (Table 1, Table S8-9) are enriched for many biological features relevant to cancer, and appear promising candidates for further investigation.

Compared with previous methods for detecting driver genes, a key feature of driverMAPS is that it models mutation rates at the base-pair level. This allows us to explicitly model how selection strength varies based on site-level functional annotations, e.g. conservation and loss-of-function status. This model-based approach can be thought of as a powerful extension of methods that detect driver genes by testing for an excess of non-synonymous vs synonymous somatic mutations (Nik-Zainal *et al*^37^, Martincorena *et al*^16^), similar to the dN/dS test in comparative genomics. Indeed, the stripped-down version of driverMAPS that uses no functional annotation or spatial model is conceptually a dN/dS test (driverMAPS-basic in Figure 4). The full version of driverMAPS, by incorporating additional functional annotations and spatial modeling, allows that some non-synonymous mutations may be more informative than others in identifying driver genes. Furthermore, by estimating parameters in a single integrated model, our approach learns how to weigh and combine the many different sources of information. The results in Figures 3 and 4 demonstrate the increased power that comes from these extensions.

Our statistical and experimental results for the mRNA methyltransferase METTL3 add to the growing evidence of links between mRNA methylation and cancer. Indeed, a recent study in myeloid leukemia cell lines^38^, found that depletion of METTL3 also leads to a cancer-related phenotype. And extensive functional studies of METTL14 in uterine cancer (Chuan He, to appear) support a role for this gene in cancer etiology. However, intriguingly, our results on METTL3 in bladder cancer, and the METTL14 results in uterine cancer suggest that they act as tumor suppressor genes, whereas the data on METTL3 in myeloid leukemia cell lines are more consistent with an oncogenic role, with depletion inducing cell differentiation and apoptosis^38^. Further studies in multiple tumor types therefore seem necessary to properly characterize the role of mRNA methylation in cancer.

Although our model incorporates many features not considered by existing methods, it would likely benefit from incorporating still more features. For example, it may be useful to incorporate data on protein structure, which affects the functional importance of amino acid residues. Further, whereas we currently use the same mutation model for all individuals, it could be helpful to incorporate individual-specific effects such as smoking-induced mutational signatures. Finally, it could be useful to extend the model to incorporate information on non-coding variation, which has been shown to be important for many human diseases including cancer. Although identifying functional non-coding variation remains a major general challenge, extending our model to incorporate features from studies of epigenetic factors such as methylation or open chromatin, has the potential to detect novel driver genes affected by non-coding somatic mutations.

## Acknowledgements

We thank Wei Du for helpful discussion and Megan McNerney for critical comments of the manuscript. We thank Peter Carbonetto for helping with software development. We also thank members of Stephens’ and He’s labs for many feedbacks during the course of this project. This work was supported by National Institutes of Health (NIH) grants MH110531 (to XH) and HG002585 (to MS).

## Code availability

driverMAPS software and procedures to reproduce the results reported in the paper can be accessed through the software website: https://szhao06.bitbucket.io/driverMAPS-documentation/docs/index.html.

## Data availability

The filtered somatic mutation lists from 20 tumor types that used as input files for driverMAPS and other comparator software are available in Zenodo (DOI: 10.5281/zenodo.1209411)^39^.

## Methods

### Data preparation

We downloaded somatic single-nucleotide mutations identified in whole exome sequencing (WES) studies for 20 tumor types from TCGA GDAC Firehose (https://gdac.broadinstitute.org/). We obtained the MAF files using firehose_get (version 0.4.6) (https://confluence.broadinstitute.org/display/GDAC/Download) and extracted position and nucleotide change information for all single-nucleotide somatic mutations. See Supplementary notes for the 20 tumor types and abbreviations.

We excluded mutations from hypermutated tumors as they likely reflect distinct underlying mutational processes. We also performed extensive filtering to exclude likely false positive mutations. For each tumor type we then generated a mutation count file that contains mutation counts (aggregated across all individuals in the tumor cohort) of all possible mutations at all sufficiently sequenced positions (see Supplementary notes). For a tumor type with 30 million bases sequenced this produces 90 million possible mutations in the mutation count file (since each nucleotide can mutate to 3 other nucleotides). The majority of counts for these possible mutations are 0s.

For each possible mutation, we annotated it with type and gene information, mutational features and functional features. We defined 9 mutation types based on nucleotide change type (such as A>T, G>A, *etc*) and genomic context (such as if inside CpG) (see Supplementary notes for definitions). We categorized mutations as Synonymous (S) or non-synonymous (NS) as described in “parameter estimation” section below. The mutational features we used include gene expression, replication timing and HiC sequencing downloaded from http://archive.broadinstitute.org/cancer/cga/mutsig. We selected 5 functional features describing mutation impact. See Supplementary notes for feature details. The features were added to the mutation count file by ANNOVAR^40^.

### Model description

We model each tumor type separately, so here we describe the model for a single tumor type. Let *Y_it_* denote the number of mutations of type *t* (defined by base substitution) at sequenced position *i*, across all samples in a cohort. Let *NS* denote the set of non-synonymous mutations.

That is, *NS* is the set of pairs (*i*,*t*) such that a mutation of type *t* at sequence position *i* would be non-synonymous. (We also include synonymous mutation with a high splicing impact score in *NS*;

see Supplementary notes.) Similarly, let *S* denote the remaining (synonymous) (*i*,*t*) pairs.

#### Background Mutation model

For synonymous mutations we assume the following “background mutation model”:

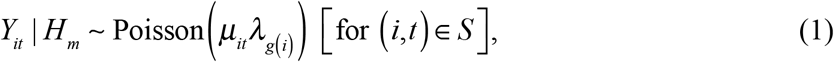

where *μ_it_* represents a background mutation rate (BMR) for mutation type *t* at position *i*, and *λ_g_*_(*i*)_ represents a gene-specific effect for the gene *g (i*) that contains sequence position *i*. Note that the parameters of this BMM do not depend on the model *m*, so 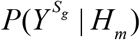 is the same for all *m*.

We allow the BMRs to depend on mutational features (e.g. the expression level of the gene) using a log-linear model:

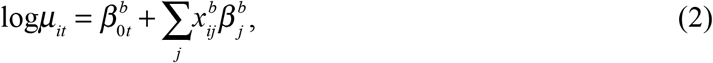

where 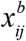 denotes the *j* -th background feature of position *i* (not dependent on mutation type), 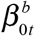 controls the baseline mutation rate of type *t*, and 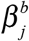 is the coefficient of the *j* -th feature. The values 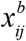 are observed, and the parameters *β ^b^* are to be estimated. To indicate the dependence of *μ_it_* on parameters *β ^b^* we write 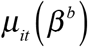.

We assume that the gene-specific effects *λ_g_* have a gamma distribution across genes:

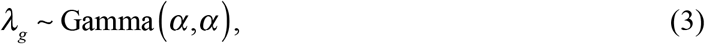

where *α* is a hyperparameter to be estimated.

#### Selection Mutation model

For non-synonymous mutations we introduce additional model-specific parameters: 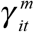 representing a selection effect (SE) for mutation type *t* at position *i* under model *m* and 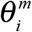 representing a spatial effect for position *i* under model *m*:

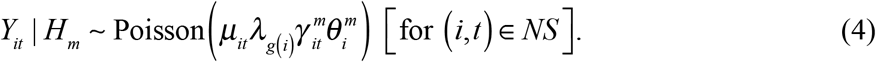

For the null model, *H*_0_, we assume no selection or spatial effect: 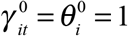.

For other models, *m* = *OG*,*TSG*, we allow the selection effect to depend on functional features (e.g. the assessed deleteriousness of the mutation), using a log-linear model:

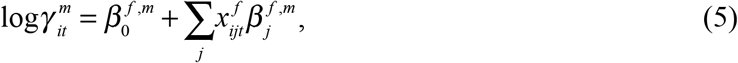

where 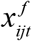 denotes the *j* -th functional feature of position *i* (this depends on mutation type; e.g. at the same position, some mutations may be more deleterious than others), 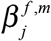 is the coefficient of the *j* -th functional feature and the intercept 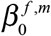 captures the overall change of mutation rate at NS sites regardless of functional impact. To indicate the dependence of 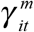 on parameters *β^f^,^m^* we write 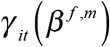.

To model the spatial effects, we use a Hidden Markov Model (HMM) with parameters Θ*^m^*,

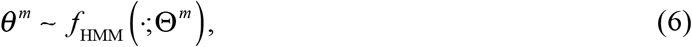

In brief, this HMM allows for the presence of mutation “hotspots” -- contiguous base-pairs with a higher rate of mutation -- and the parameters include the average hotspot length and intensity of selection (ρ). See Supplementary note for details.

### Parameter estimation

#### Background mutation model

To simplify inference we took a sequential approach to parameter estimation. First we infer parameters *β ^b^*,*α* of the BMM using the synonymous mutation data at all genes. Let *S_g_* denote the subset of synonymous mutations *S* in gene *g*, and 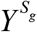 denote the corresponding observed counts:

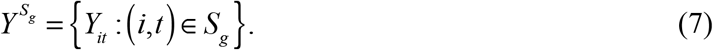

Based on the synonymous mutation data, the likelihood for gene *g* is:

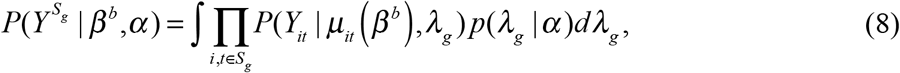

which has a closed form (see Supplementary note). Assuming independence across genes yields the likelihood for synonymous mutations:

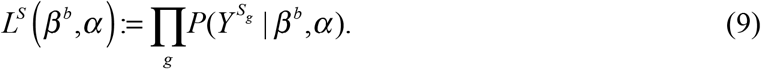

We maximize this likelihood, using numerical optimization, to obtain estimates 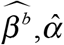 for *β ^b^*,*α*.

By ignoring the non-synonymous mutation data when fitting the BMM we may lose some efficiency in principle, but we gain considerable simplification in practice.

#### Selection mutation model

We next estimate the model-specific parameters *β ^f,m^*. For *m* = *OG*,*TSG*. During this step we ignore the HMM model (i.e. we set 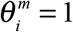), motivated by the fact that spatially-clustered mutations are relatively rare and so should not significantly impact the estimates of *β ^f,m^* For *m* = *OG* we estimate *β ^f^* ^,*m*^ using the non-synonymous mutation data from a curated list *G_OG_* of 53 OGs. Estimation for *β^f,TSG^* is identical except that we replace this list with a curated list *G_TSG_* of 71 TSGs. Let *G_m_* denote these sets of training genes. Let 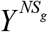 denote the counts of non-synonymous mutations in gene *g*.

Assuming independence across genes, the likelihood for *β^f,m^* is:

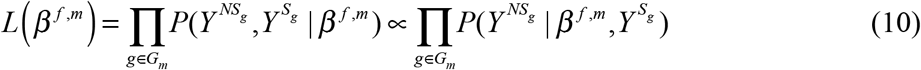

where the second line follows because 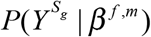 does not depend on *β^f,m^*. The term in this likelihood for gene *g* is given by:

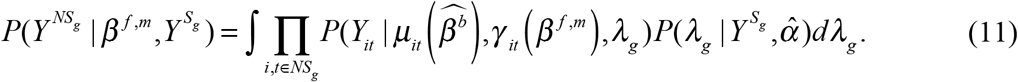

It can be shown that

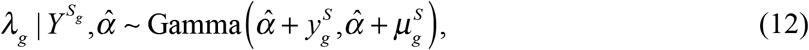

where 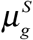 and 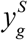 are, respectively, the expected (considering only mutational features) and observed number of synonymous mutations in gene *g* (see Supplementary notes). The conditional mean of this distribution is 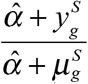, so if 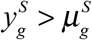, then 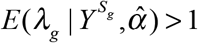.

We obtained the MLE of *β^f,m^* by maximizing the likelihood (Equation 10) numerically, and obtain corresponding estimated standard errors using the curvature of the likelihood (see Supplementary notes). In tumor types with low mutation rates or sample sizes, these standard errors can be relatively large, so we borrow information from other tumor types to ‘’stabilize’’ these estimates. Specifically we use the adaptive shrinkage method^20^ to “shrink” estimated values of *β^f^* ^,*m*^ in each tumor type towards the mean across all tumor types. This shrinkage effect is strongest for tumor types with large standard errors (Figure S7).

#### HMM parameters

Having estimated *β^b^*,*α* and *β ^f,m^*, we fix their values and estimate the HMM parameters Θ*^m^* for *m* = *OG*,*TSG*. The likelihood function involves marginalization of the hidden states of the Markov chain, which can be performed efficiently using standard methods for HMMs. We estimate Θ*^m^* by maximizing this likelihood numerically. See Supplementary note for details.

### Gene classification

Having estimated the model parameters as above, for each gene *g*, we compute its Bayes Factor for being a driver gene as:

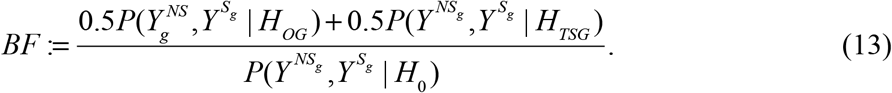

The equal weights in the numerator of this BF assume that OGs and TSGs are equally common. This BF simplifies to

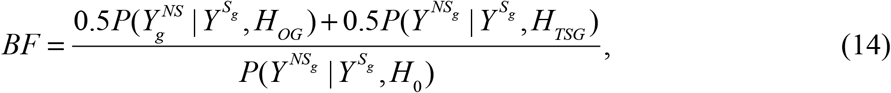

because 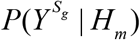 is the same for every *m*. Computing the terms 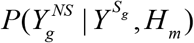 is performed using (Equation 11) above, substituting the estimated model parameters for each model *m* (see Supplementary notes).

After obtaining the BFs, we can compute the posterior probability of being a driver gene (either *OG* or *TSG*) for every gene, and estimate the Bayesian FDR^41^ for any given BF threshold. This step requires estimation of the proportion of driver genes, which we do by maximum likelihood (see Supplementary notes).

### Simulations

For power analysis shown in Figure 3(a), we randomly picked a gene (*ERBB3*) and for a given number of samples, we simulated mutations under positive selection and assessed the power of detecting this gene as positively selected using different methods. We simulate synonymous mutations at predefined background mutation rates (BMRs); we simulate positively selected mutations at elevated mutation rates for nonsynonymous sites and hotspot sites (generated by a Markov chain). This simulation procedure was performed many times and each time we obtained *p* value for each method. Power is defined as the fraction of simulations with significant *p* values (*p* < 0.05). The test statistics for “dN/dS” method is the likelihood ratio of between Poisson models under elevated mutation rates and BMRs. The test statistics for “cluster” method is the maximum number of mutations within 3bp windows normalized by overall mutation rates. Null distributions of test statistics are obtained by simulations with mutation rates for all sites equal to BMRs. *p* value for “combined” method is obtained by combining *p* values of “dN/dS” and “cluster” using Fisher’s method.

For simulation performed in Figure 3(b) and (c), we simulated positively selected mutations for 324 genes and neutral mutations for the rest genes. 124 out of the 324 genes are known TSGs or OGs, the same as the training set for driverMAPS. The rest 200 genes were randomly sampled from all genes. The 71 TSGs used for training and 120 out of the 200 randomly sampled genes were simulated under H_TSG_. The 53 OGs used for training and 80 out of the 200 randomly sampled genes were simulated under H_OG_. For neural genes and synonymous sites in positively selected genes, we simulated mutations at predefined BMRs; for nonsynonymous in positively selected genes, we simulated mutations at increased rates based on its functional annotations and hotspot status generated by Markov Chain. We removed the 124 genes used as the training set for driverMAPS from results in all methods and only the rest 200 genes were used as the true set for the ROC curve to ensure fair comparisons.

For all simulations, the predefined BMRs, effect sizes for functional annotations and spatial clustering hotspot rated parameters were estimates by driverMAPS using UCS data (Table S1-S5, UCS parameters). We re-estimated these parameters when running driverMAPS.

### Comparison of gene prediction results from different methods

When comparing methods, we used the same mutation data (after filtering) and the same nominal FDR threshold (0.1) for each method. Because driverMAPS used 124 known cancer genes as a training set, to avoid bias towards this subset of genes when computing precision or power for driverMAPS, we ran MAPs using a leave-one-out strategy. We perform 124 runs, each time omitting one TSG/OG from the training set and estimating model parameters from the remaining genes, and then count the omitted gene as “significant” only if this TSG/OG is significant (FDR<0.1) in this run. We then calculate precision as the percentage of significant known cancer genes of all significant genes. All data related to driverMAPS (basic, +feature and full version) presented in Figure 3 were obtained in this way. In fact, estimated model parameters are quite stable across runs, and so the overall result is similar to a single run not using this “leave-one-out” strategy.

### Cell lines, siRNA knockdown and plasmid transfection

The T24 cells used in this study were purchased from ATCC (HTB-4) and grown in McCoy’s 5A medium (Gibco, 16600) supplemented with 10% FBS (Gibco), and 1% Penicillin-Streptomycin (Gibco, 15140). The 5637 cells used in this study were purchased from ATCC (HTB-9) and grown in RPMI-1640 medium (Gibco, 11875) supplemented with 10% FBS and 1% Penicillin-Streptomycin. Construction of the pcDNA3 plasmids for the expression of METTL3 in mammalian cells was described previously. All siRNAs were ordered from QIAGEN. Allstars negative control siRNA (1027281) was used as siRNA control. Sequences METTL3 is 5’- CGTCAGTATCTTGGGCAAGTT-3’. Transfection was achieved by using Lipofectamine RNAiMAX (Invitrogen) for siRNA, or Lipofectamine 2000 (Invitrogen) for the plasmids following manufacturer’s protocols.

### *In vitro* assay for m^6^A methyltransferase activity

The recombinant, His-tagged proteins METTL14 with wildtype or mutant METTL3 were expressed in 1 LB Ecoli expression system and purified through Ni-NTA affinity column according to a previously published procedure^42^. Protein purity was assessed by SDS-PAGE, and protein concentration was determined by UV absorbance at 280 nm. We performed an *in vitro* methyltransferase activity assay in a 50 *μ*L reaction mixture containing the following components: 0.15 nmol RNA probe, 0.15 nmol each fresh recombinant protein (METTL14 combination with an equimolar ratio of METTL3 or mutant METTL3), 0.8 mM *d3*-SAM, 80 mM KCl, 1.5 mM MgCl_2_, 0.2 U *μ* L-1 RNasin, 10 mM DTT, 4% glycerol and 15 mM HEPES (pH 7.9). The reaction was incubated for 12 h at 16 °C, RNA was recovered by phenol/chloroform (low pH) extraction followed by ethanol precipitation and was digested by nuclease P1 and alkaline phosphatase for LC-MS/MS detection. The nucleosides were quantified by using the nucleoside-to-base ion mass transitions of 285 to 153 (*d3*-m^6^A) and 284 to 152 (G).

### RNA isolation

Total RNA was isolated with TRIZOL reagent (Invitrogen). mRNA was extracted from the total RNA using the Dynabeads^®^ mRNA Purification Kit (Invitrogen), followed by removal of contaminating rRNA with the RiboMinus transcriptome isolation kit (Invitrogen). mRNA concentration was measured by UV absorbance at 260 nm.

### LC-MS/MS quantification of m^6^A in poly(A)-mRNA

100-200 ng of mRNA was digested by nuclease P1 (2 U) in 25 *μ* L of buffer containing 25 mM of NaCl, and 2.5 mM of ZnCl_2_ at 42 °C for 2 h, followed by the addition of NH_4_HCO_3_ (1 M, 3*μ* L) and alkaline phosphatase (0.5 U) and incubation at 37 °C for 2 h. The sample was then filtered (0.22 m pore size, 4 mm diameter, Millipore), and 5 *μ*L of the solution was injected into the LC-MS/MS. The nucleosides were separated by reverse phase ultra-performance liquid chromatography on a C18 column with online mass spectrometry detection using Agilent 6410 QQQ triple-quadrupole LC mass spectrometer in positive electrospray ionization mode. The nucleosides were quantified by using the nucleoside to base ion mass transitions of 282 to 150 (m^6^A), and 268 to 136 (A). Quantification was performed by comparison with a standard curve obtained from pure nucleoside standards run with the same batch of samples. The ratio of m^6^A to A was calculated based on the calibrated concentrations.

### Cell proliferation assay

5000 cells were seeded per well in a 96-well plate. The cell proliferation was assessed by assaying the cells at various time points using the CellTiter 96^®^ Aqueous One Solution Cell Proliferation Assay (Promega) following the manufacturer’s protocols. For each cell line tested, the signal from the MTS assay was normalized to the value observed ~24 hours after seeding.

